# High dynamic range shortwave infrared (SWIR) imaging of mice with an InGaAs camera

**DOI:** 10.1101/2025.11.06.687088

**Authors:** Amish Patel, Xingjian Zhong, Mallory Moffett, Yidan Sun, Allison M. Dennis

## Abstract

**Significance:** While shortwave infrared (SWIR) imaging provides superior tissue penetration and reduced autofluorescence for preclinical applications, quantitative fluorescence analysis is hindered by the limited dynamic range of InGaAs cameras, forcing a focus on either bright or dim anatomical features.

**Aim:** We develop a high dynamic range (HDR) imaging method specifically adapted for the high-noise characteristics of InGaAs detectors to enable quantitative fluorescence imaging across wide intensity ranges. We demonstrate that one-time camera calibration based on a series of images encompassing the range of radiance intensities enables all subsequent image processing.

**Approach:** We modified classical HDR algorithms with exposure-time-dependent dark current subtraction, preprocessing to exclude saturated and noisy pixels before camera response function recovery, and dynamic weighting range adjustment to account for shrinking intensity ranges at longer exposures. HDR image processing effects on preclinical imaging outcomes were analyzed using indocyanine green and SWIR-emitting PbS/CdS quantum dots in mouse models.

**Results:** HDR imaging achieved a 22 dB improvement in dynamic range over single exposures, enabling simultaneous quantification across more than three orders of magnitude of fluorophore concentration. *In vivo* studies showed improvements in contrast-to-noise ratios across all anatomical features, with improvements in vascular contrast while maintaining quantitative accuracy. After one-time camera calibrations, this approach enables rapid processing of subsequent datasets.

**Conclusions:** This software-based HDR SWIR imaging approach eliminates exposure parameter optimization and enables comprehensive biodistribution analysis across all anatomical structures from a single acquisition sequence, significantly streamlining preclinical imaging workflows while preserving quantitative accuracy.

## 1 Introduction

Fluorescence imaging has emerged as an indispensable tool for molecular imaging, offering high spatial resolution and excellent signal-to-noise ratios (SNR) without the risks associated with ionizing radiation.^1,2^ However, the absorption and scattering of biological tissues severely limits imaging depth when using conventional fluorophores and silicon-based detectors. While near infrared (NIR-I) fluorophores operating in the 700-900 nm window have partially addressed these limitations,^3–5^ persistent tissue autofluorescence and photon scattering continue to hinder visualization of deep anatomical structures and organs.^6–8^

The shortwave infrared (SWIR) imaging window (1000-1700 nm, a.k.a., NIR-II) offers compelling advantages over NIR-I imaging, including dramatically reduced tissue autofluorescence^9,10^ and decreased photon scattering.^11^ These optical improvements translate directly into superior image contrast for deep tissue features, enabling visualization of organs and vascular structures previously inaccessible to non-invasive fluorescence imaging.^12–14^ Despite advances in SWIR detector technology,^15–17^ the limited dynamic range of commercial InGaAs cameras presents end-users with a challenging choice: use exposure settings that capture dim features but result in signal saturation in high-uptake organs like the liver and spleen or non-saturating exposure times with which dimmer features disappear into detector noise. For the majority of SWIR imaging users who cannot modify their camera hardware, this fundamental limitation prevents simultaneous quantification across the orders-of-magnitude concentration and intensity differences encountered in biomedical fluorescence applications, forcing researchers to sacrifice either their brightest or dimmest signals and hindering quantitative analysis.

While high dynamic range (HDR) imaging offers a potential solution by combining multiple exposures to extend the measurable intensity range, classic HDR methods developed for silicon cameras are inadequate for InGaAs SWIR detectors. Established algorithms assume negligible detector noise, which is reasonable for silicon-based cameras but inappropriate for InGaAs SWIR detectors.^15^ Previous examples of SWIR HDR imaging have focused exclusively on reflected light applications in machine vision and remote sensing, and choose intensity values from images taken at multiple exposure times without recovering the camera response function or establishing proper noise models.^16,17^ Notably, no validated HDR method has been specifically designed for the high-noise characteristics and exposure time-dependent dark current (DC) behavior of InGaAs cameras for use in biomedical fluorescence imaging applications.

We address this critical gap by developing a practical HDR imaging approach that works with any existing InGaAs SWIR camera system, requiring no hardware modifications or proprietary components. Our method adapts a classic HDR algorithm,^18^ introducing three key modifications that address the unique challenges of SWIR detectors: *(1)* a one-time characterization of exposure-time-dependent DC that can be applied to all subsequent imaging, *(2)* preprocessing to exclude saturated and noisy pixels before CRF recovery, and *(3)* dynamic weighting adjustment to account for the shrinking usable signal window at longer exposures. After initial calibration, which requires only a series of DC and reflectance or fluorescence images, these models enable rapid HDR processing of any SWIR fluorescence dataset acquired with the same camera. This approach enables > 20 dB improvement in dynamic range using only software processing and standard imaging protocols, eliminating cost and complexity barriers. This advance is particularly significant for preclinical imaging applications, where fluorescent contrast agents accumulate at vastly different concentrations across anatomical structures. By enabling quantification across more than three orders of magnitude of fluorophore concentration from a single imaging sequence, our HDR approach eliminates the need for prior knowledge of optimal exposure parameters and streamlines experimental workflows while preserving quantitative accuracy.

We demonstrate that these adaptations successfully recover accurate camera response functions despite high detector noise, enabling generation of radiometrically calibrated images across wide intensity and fluorophore concentration ranges. We systematically validate our approach from camera characterization through *in vitro* standards and *in vivo* imaging to show that HDR SWIR imaging can enable signal quantification in preclinical imaging applications. Our results demonstrate that this accessible HDR method enables quantitative analysis across all anatomical structures from a single acquisition sequence, providing the quantitative imaging capabilities previously available only through expensive, specialized hardware solutions.

## 2 Methods

### 2.1 Image acquisition

SWIR fluorescence images were acquired using an IR VIVO multispectral preclinical imaging system (Photon etc., Montreal, Canada) equipped with a ZephIR™ 1.7 InGaAs camera (640 × 512 pixels, 15 µm pixel pitch, spectral range 500-1630 nm). The system uses thermoelectric cooling to maintain stable detector temperature at -80°C throughout imaging sessions. Three excitation lasers (670, 760, and 808 nm) provide homogeneous rectangular illumination across the field of view. For this study, we used 808 nm excitation at power densities of 0.5-2 mW/mm^2^ (adjustable based on working distance).

Emission was collected through various filters mounted in an automated filter wheel: NIR-II LP (1000-1600 nm), 1250 LP (1250-1600 nm), and bandpass filters (50 nm bandwidth) centered at 1150, 1200, 1300, 1350, 1500, and 1550 nm. For HDR imaging, we acquired 4-8 frames in high gain mode at increasing exposure times, typically in a geometric series ranging from 0.01 to 6.4 s with 2-fold increments. All image processing was performed using custom algorithms written in Python with visualization performed using Python (matplotlib version 3.8.4), GraphPad Prism (version 10.0), and Fiji (ImageJ version 1.53t).

### 2.2 HDR image processing workflow

Our HDR imaging protocol consists of two distinct phases: an initial calibration phase and subsequent imaging and image processing phases (Fig. 1). The calibration phase, performed once for the imaging system, involves comprehensive DC characterization across the full range of exposure times and recovery of the camera response function (CRF) using images representing the full range of pixel intensities generated with reflectance standards or fluorescent datasets. This calibration yields a per-pixel DC model that predicts noise for any exposure time, a per-pixel saturation value (*S_max_*), and a CRF *g*(*z*) that maps pixel values to irradiance, i.e., exposure. For routine preclinical imaging, we apply these pre-established calibration models to efficiently process new datasets. Each HDR acquisition involves collecting 2-8 frames at varying exposure times, applying DC correction using our calibrated model, capping the saturated pixel intensities, and fusing the frames using the CRF, emphasizing the most reliable pixel intensity ranges using the weighting function.

**Figure 1.**
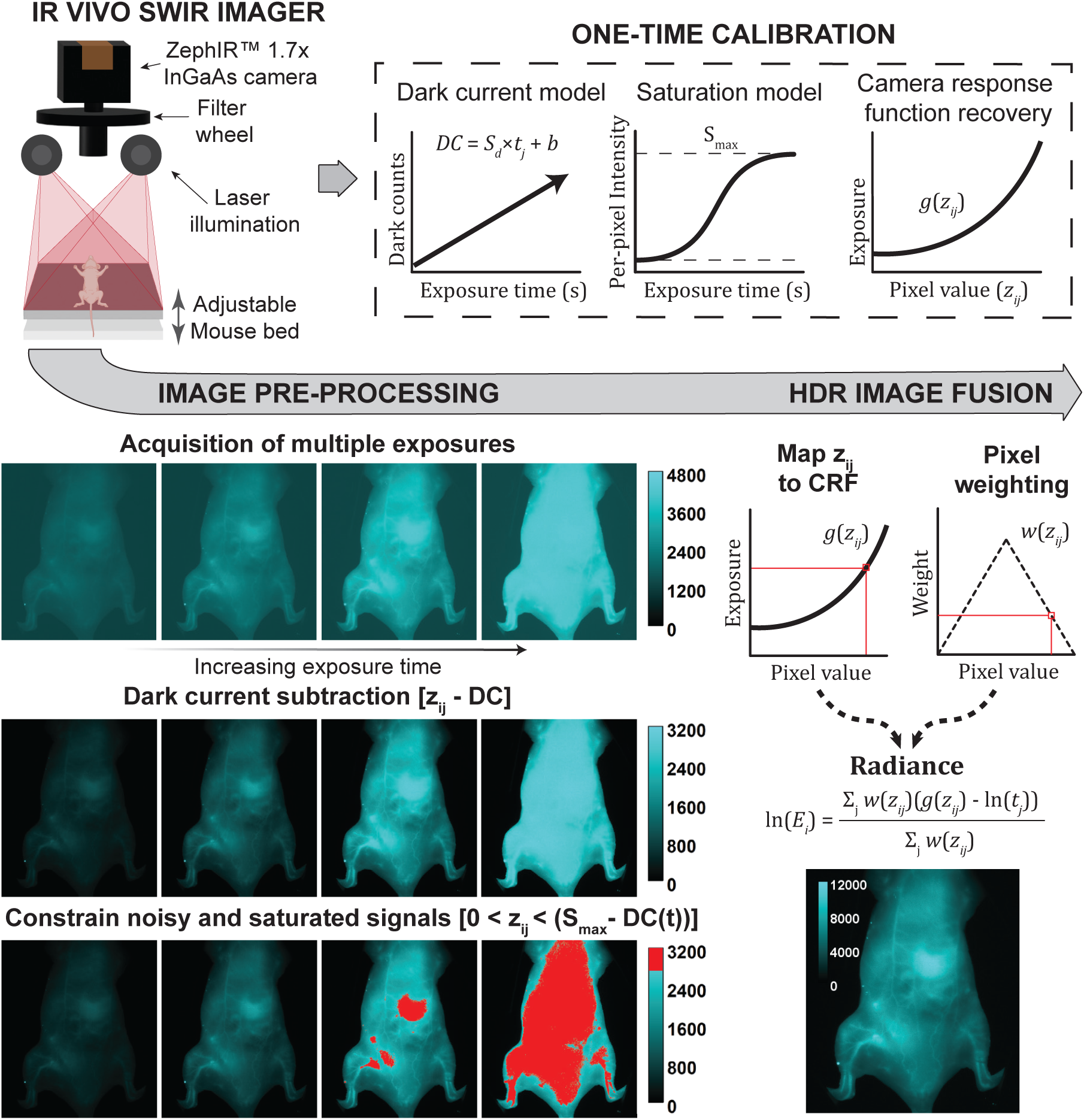
SWIR imaging system and HDR processing workflow. Schematic of the IR VIVO preclinical imager: laser excitation modules (670, 760, 808 nm) provide homogenous rectangular illumination across the field of view with a maximum illumination power of 2 mW/mm^2^; fluorescence emission from samples placed on the imaging stage passes through a SWIR lens and a filter wheel before reaching a thermoelectrically cooled InGaAs detector. Each image is denoised by DC subtraction and constrained to cap the saturated pixel intensities prior to HDR image processing to improve final image accuracy. Following pre-processing, each pixel value 𝑧_*ij*_ per position *i* and time *j* is mapped to a previously recovered camera response function 𝑔(𝑧_*ij*_) to quantify irradiance at the camera sensor. A weighting function 𝑤(𝑧_*ij*_) is applied to the exposure values to emphasize data in the more reliable pixel intensity range away from the extremes when producing the HDR radiance map. The dark current noise model and camera response function required to perform preprocessing and HDR image fusion, respectively, are generated in a one-time sensor calibration step, producing the parameters used in all subsequent image processing.

### 2.3 Dark current (DC) and saturation characterization

#### DC characterization

We acquired 100 dark frames (camera shutter closed) at exposures ranging from 0.001 to 32 s. These dark frames were averaged to reduce the impact of thermal noise and extract camera bias. Following methods adapted from Shaikh et al.,^15^ we modeled the DC noise as a function of exposure time (*t*) and pixel position (*x, y*) using these averaged dark frames:

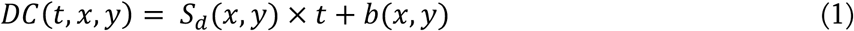

where 𝑆_*d*_ is the DC rate (counts/s), 𝑡 is exposure time (s), and 𝑏 represents the combined contribution of camera bias, which includes read noise and fixed pattern noise. We determined 𝑆_*d*_ and 𝑏 for each pixel through linear regression. This model defines the effective noise floor 𝑍_*min*_ as the DC(t) for each pixel at a given exposure time, which is considered the lower bound of pixel values z during HDR image processing.

#### Saturation characterization

To determine the maximum pixel value (𝑆_*max*_), we illuminated a Lambertian reflectance standard (Teflon sheet) using a broadband calibration light source (Ocean Optics) positioned at ∼20° to minimize specular reflection. We acquired images at exposure times from 0.001 to 30 s and fitted each pixel response to a sigmoid function:

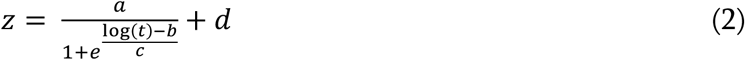

where *a* represents the response magnitude, *b* the temporal offset, *c* the transition steepness, and *d* the baseline offset of pixel intensity *z*. The pixel saturation point 𝑆_*max*_ was defined as the sum (*a* + *d*), representing the asymptotic maximum for each pixel in raw data acquisition. This value establishes 𝑍_*max*_, the upper limit for pixel intensities *z*, beyond which pixel response becomes unpredictable and must be excluded from HDR processing. 𝑍_*max*_ is further adjusted according to data preprocessing to dynamically define the bounds of *z*.

### 2.4 Camera Response Function Recovery

Digital cameras exhibit nonlinear relationships between incident irradiance and reported pixel values. To generate radiometrically accurate HDR images, we must first characterize this camera response function (CRF). We adapted the method of Debevec and Malik to recover the CRF by minimizing:^18^

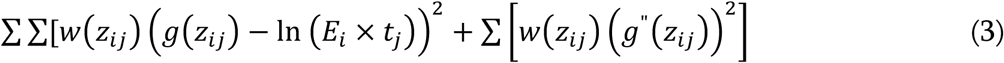

where *g*(*z_ij_*) is the response function mapping pixel intensity *z_ij_* to log irradiance, 𝐸_*i*_ is the irradiance at pixel with spatial location *i*, 𝑡_*j*_ is exposure time with exposure number *j*, 𝑤(*z_ij_*) is the weighting function, and the second term enforces smoothness.

To collect CRF calibration data, we imaged a checkerboard pattern under controlled illumination at 6 exposure times between 0.01 and 0.32 s. The varied reflectance of the checkerboard ensures sufficient representative sampling across the camera response range. We randomly sampled 5,000 pixels, yielding 30,000 pixel intensity values across the 6 exposure times, then excluded up to half of the pixels if one or more of their exposures were outside the bounds of 𝑍_*min*_ and 𝑍_*max*_ (i.e., removed pixels with under-or oversaturated intensities at any exposure). We verified the system was sufficiently overdetermined with 𝑁(𝑃 − 1) > (𝑍_*max*_ − 𝑍_*min*_), where 𝑁 is the number of sampled pixels and 𝑃 is the number of exposures. The recovered CRF was validated to be monotonically increasing, confirming proper convergence. Fluorescence images may be used for CRF recovery, though reflectance-based samples provide a higher number of bright pixels and thus may require fewer samples.

### 2.5 Pixel Weighting for High-Noise SWIR Imaging

A weighting function is used to emphasize more reliable pixel values in the CRF recovery and HDR fusion processes. Debevec’s triangular weighting function 𝑤(𝑧) minimizes pixels near 𝑍_*min*_ and 𝑍_*max*_, while pixels near the average of the two extremes 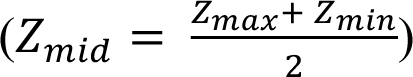 are weighted close to their full value:^18^

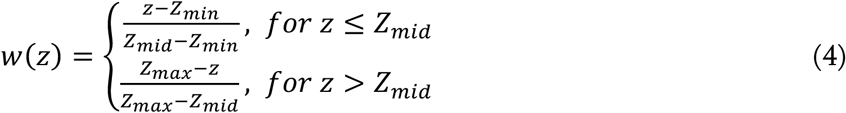

Unless noted otherwise, we subtract the DC from each frame prior to applying the weights, effectively remapping 𝑍_*min*_ to 0 and adjusting 𝑍_*max*_ accordingly.

### 2.6 HDR Image Fusion

We fuse multiple exposures into a single radiometrically calibrated HDR image using the CRF generated in section 2.4. We calculate the log radiance at each pixel using:

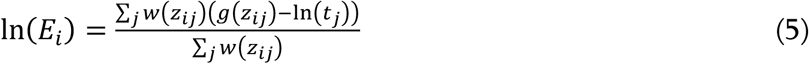

where 𝐸_*i*_ is the radiance at pixel *i*, *g*(*z_ij_*) maps pixel values to log irradiance using the recovered CRF, *t_j_* is the exposure time, and 𝑤(*z_ij_*) applies the triangular weighting function described in section 2.5. The resulting radiance map provides relative irradiance values that can be quantitatively compared across images acquired with the same camera and CRF.^18^

### 2.7 SWIR Contrast Agents

We evaluated our HDR method using two (pre)clinically relevant SWIR fluorophores. Absorption spectra were measured using a Jasco® V-780 UV-Visible/NIR spectrophotometer, and photoluminescence spectra were measured using the IR VIVO imager in hyperspectral mode (Figure 2). Indocyanine-green (ICG; Cayman Chemical) was dissolved in DMSO and diluted in sterile saline for injection. PbS/CdS core/shell quantum dots were synthesized as previously described and transferred to aqueous media using micelle encapsulation.^14,19^ Optical spectra of the PbS/CdS QDs were measured with the QDs suspended in tetrachloroethylene (TCE) to avoid the impact of water absorption.

**Figure 2.**
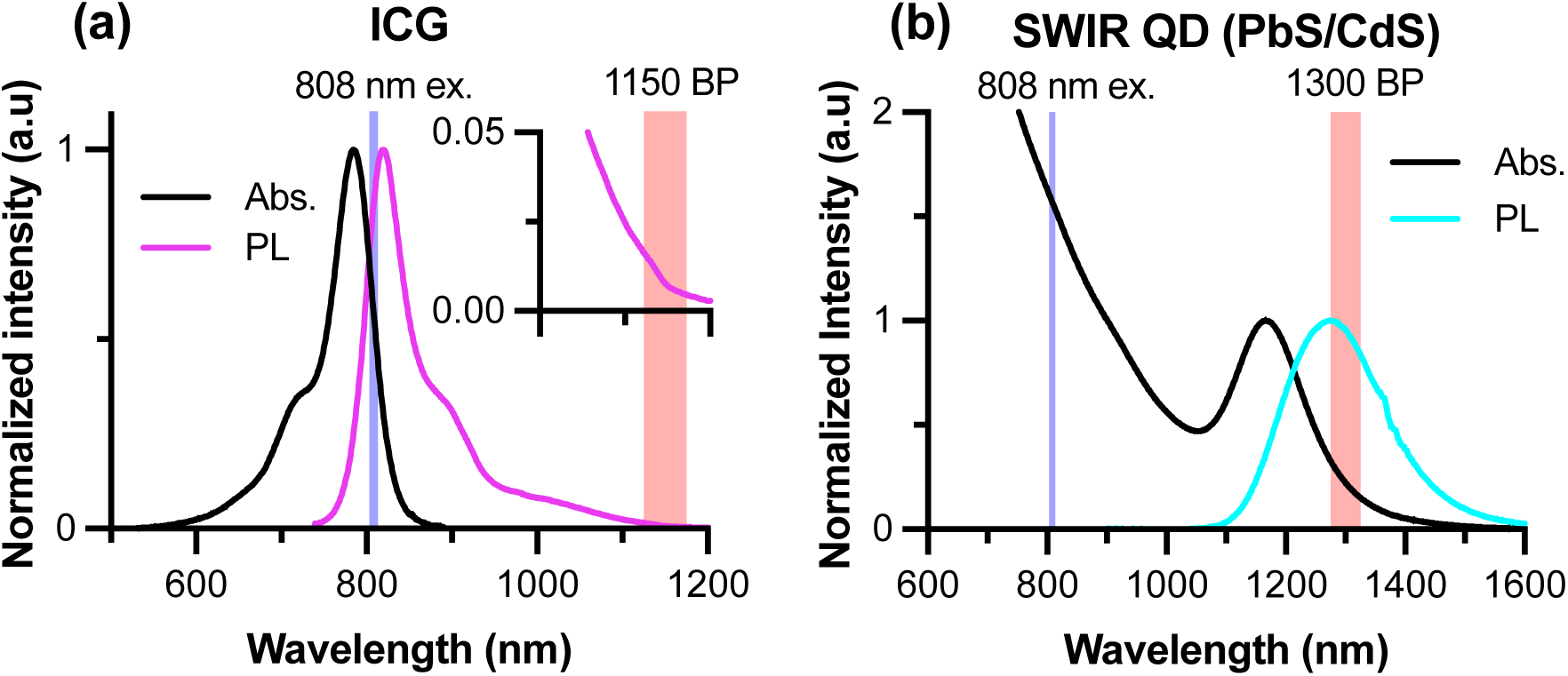
Normalized absorption and photoluminescence (PL) spectra of SWIR contrast agents. (a) ICG spectra with the 808 nm excitation laser and 1150 nm bandpass filter used in subsequent imaging indicated by the light blue and red shaded regions, respectively. (b) Absorbance and emission spectra of SWIR QDs suspended in TCE with the 808 nm excitation laser and 1300 nm bandpass filter used in subsequent imaging indicated by the light blue and red shaded regions on the plot, respectively.

### 2.8 Animal Imaging

All procedures were approved by the Northeastern University IACUC (Protocol #22-1027R). NU/J nude mice (Jackson Laboratory, strain #002019) were fed a chlorophyll-free diet (OpenStandard Diet without dye, D11112201N, Research Diets, Inc.) for at least one week prior to imaging to minimize tissue autofluorescence.^10^ Mice were anesthetized with 2-2.5% isofluorane mixed with house air at a flow rate of 1.5 L/min at least 5 min prior to and throughout imaging, and were observed for 30 min after imaging was complete to ensure healthy behavior. For biodistribution imaging, mice received retro-orbital injections of either (1) PbS/CdS quantum dots at 5 mg/kg total Pb+Cd dose (n = 4 8-week-old mice) or (2) ICG at 0.5 mg/kg (n = 4 20-week-old mice). Total injection volume ranged from 50-80 μL for PbS/CdS quantum dots and 80-100 μL for ICG. For each HDR acquisition, 4-8 frames with exposure times ranging from 0.01 to 6.4 s were collected, maintaining total light exposure within ANSI laser safety limits (<2.0 mW/mm^2^). Emission filters were chosen to optimize contrast for each fluorophore: 1300 nm bandpass for QDs and 1150 nm bandpass for ICG.

### 2.9 Quantitative and Statistical Analysis

Image analysis was performed using custom Python scripts. For specific images (scene dynamic range for single exposure or HDR images), dynamic range (DR) was calculated as:

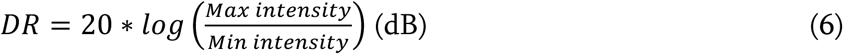

For a single exposure, the maximum potential DR was dependent on the exposure time:

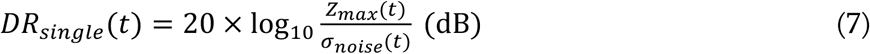

where *Z_max_* is the maximum detectable signal before saturation and 𝜎_*noise*_ (𝑡), is the standard deviation of background noise. The DR is reduced at longer single exposure times because both the lowest detectable signal *DC*(*t*) and the standard deviation of the dark counts 𝜎_*DC*_ (𝑡), which is the primary contributor to 𝜎_*noise*_ (𝑡), increase with exposure time. After DC subtraction using the empirically determined model for *DC*(*t*), the noise floor includes both read noise (*σ_read_*) and uncertainty from the DC model:

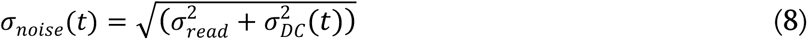

where 𝜎_*DC*_ (𝑡) represents the standard deviation in the DC model at exposure time *t*. For HDR radiance maps generated from multiple exposures, the theoretical maximum dynamic range was calculated as:

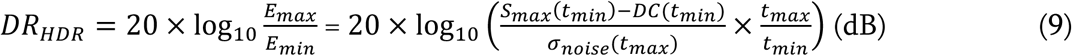

where the maximum radiance 𝐸_*max*_ over the minimum radiance 𝐸_*min*_ is the detector-limited dynamic range using the brightest signal from the shortest exposure (𝑆_*max*_ − 𝐷𝐶(𝑡) at *t_min_*) and the noise floor from the longest exposure (𝜎_*noise*_ (*t_max_*)), and the factor 𝑡_*max*_ /𝑡_*min*_ accounts for the additional dynamic range gained through exposure bracketing.

## 3 Results and Discussion

### 3.1 InGaAs Camera Characterization Defines Operating Parameters

Successful HDR imaging requires precise understanding of camera behavior across the full range of operating conditions. We first characterized the DC and saturation properties of our thermoelectrically cooled InGaAs camera to establish the usable dynamic range and inform our HDR algorithm design. The DC and saturation values were determined and then utilized in image preprocessing on a per-pixel basis, preserving the full DR of each individual pixel and ensuring that image preprocessing does not introduce noise or artifacts by treating the pixel ranges homogenously.

#### DC characterization

The InGaAs detector is thermoelectrically cooled to -80°C, eliminating temperature-dependent variations in DC and enabling stable operation across imaging sessions. This consistent cooling allows for characterization of the dark current as a function of exposure time alone, without needing to account for temperature fluctuations. Analysis of 100 dark frames acquired at each exposure time across a geometric sequence ranging from 0.001 to 32 s revealed highly predictable behavior. The linear model *DC*(*t*) = *S_d_* × *t* + *b* accurately described pixel response (R^2^ = 0.993) within the range of 0.01 – 16 s (Figure 3a). Below 0.01 s, read and shot noise dominated the signal, while above 16 s, DC approached pixel saturation and exhibited nonlinear behavior. The upper operational limit occurred at approximately 20 s, where the bias term *b* approached the saturation point, leaving no headroom for signal. The regression was performed independently for each pixel and the coefficients *S_d_* and *b* saved for rapid reconstruction of the pixel-specific DC model. Despite relatively low pixel-to-pixel variance in both slope (*S_d_* = 181.8 ± 6.8 counts/s) and bias (*b* = 1171.7 ± 6.9 counts), per-pixel calibration ensures optimal noise subtraction across the entire sensor, as even small variations in DC can impact quantitative measurements when working near the noise floor.

**Figure 3.**
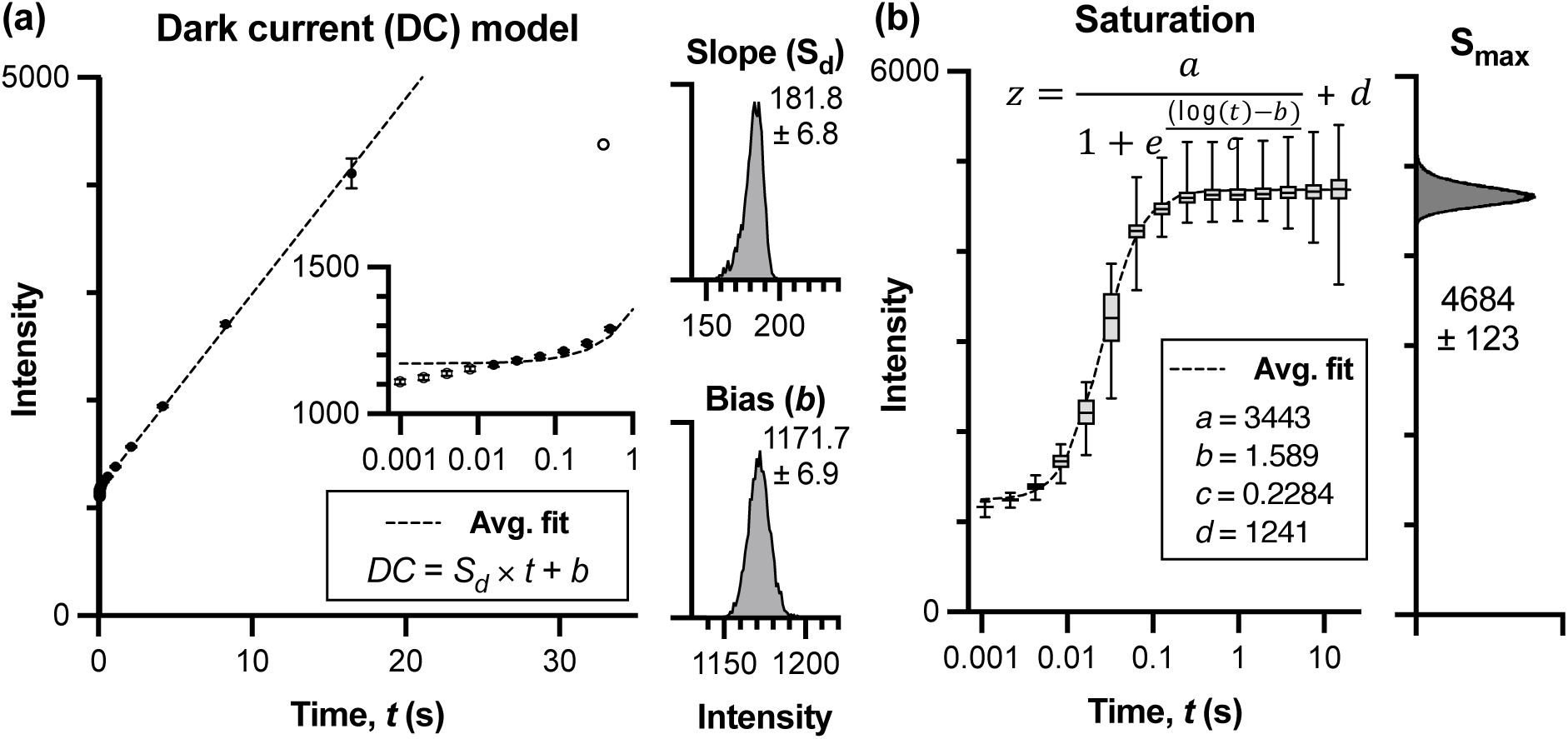
Characterization of InGaAs camera dark current (DC) and saturation behavior. (a) DC characterization using the mean pixel intensity of 100 dark frames at each exposure time over a range from 0.001 to 32 s. The linear model *DC(t)* = *Sd* × *t* + *b* accurately describes pixel behavior (R^2^ = 0.993) between 0.01 and 16 s (filled markers). Beyond this range (open markers), DC exhibits nonlinear behavior and was excluded from the DC model and subsequent experimental imaging. Inset histograms show the distribution of slope (*Sd* = 181.8 ± 6.8 counts/s) and bias (*b* = 1171.7 ± 6.9 counts) values across all pixels. (b) Saturation characterization using reflectance measurements from a Teflon standard. Pixel intensities were fitted to sigmoid functions to determine saturation points. The asymptotic maximum *Smax* (4684 ± 123 counts), calculated as a sum of the magnitude (*a*) and offset (*d*) parameters, defines the upper bound of reliable signal for each pixel. Error bars represent standard deviation across pixels.

#### Saturation characterization

Imaging a Lambertian reflectance standard across exposure times from 0.001 to 32 seconds revealed consistent saturation behavior across pixels (Fig. 3b). The sigmoid model (Equation 2) accurately captured the transition to saturation, with the asymptotic maximum *S_max_* = 4684 ± 123 counts showing relatively low variance across the sensor. We use the per-pixel *S_max_* because even small variations in saturation points could cause some pixels to saturate earlier than others, leading to inaccurate HDR fusion if not properly accounted for.

### 3.2 Camera Response Function Recovery Requires Careful Signal Preprocessing

Having established the detector operating range, we next recovered the camera response function (CRF) for radiometric calibration. Initial attempts using standard HDR algorithms failed due to the high noise characteristics of the InGaAs sensor. Figure 4 demonstrates the critical importance of signal preprocessing for accurate CRF recovery.

**Figure 4.**
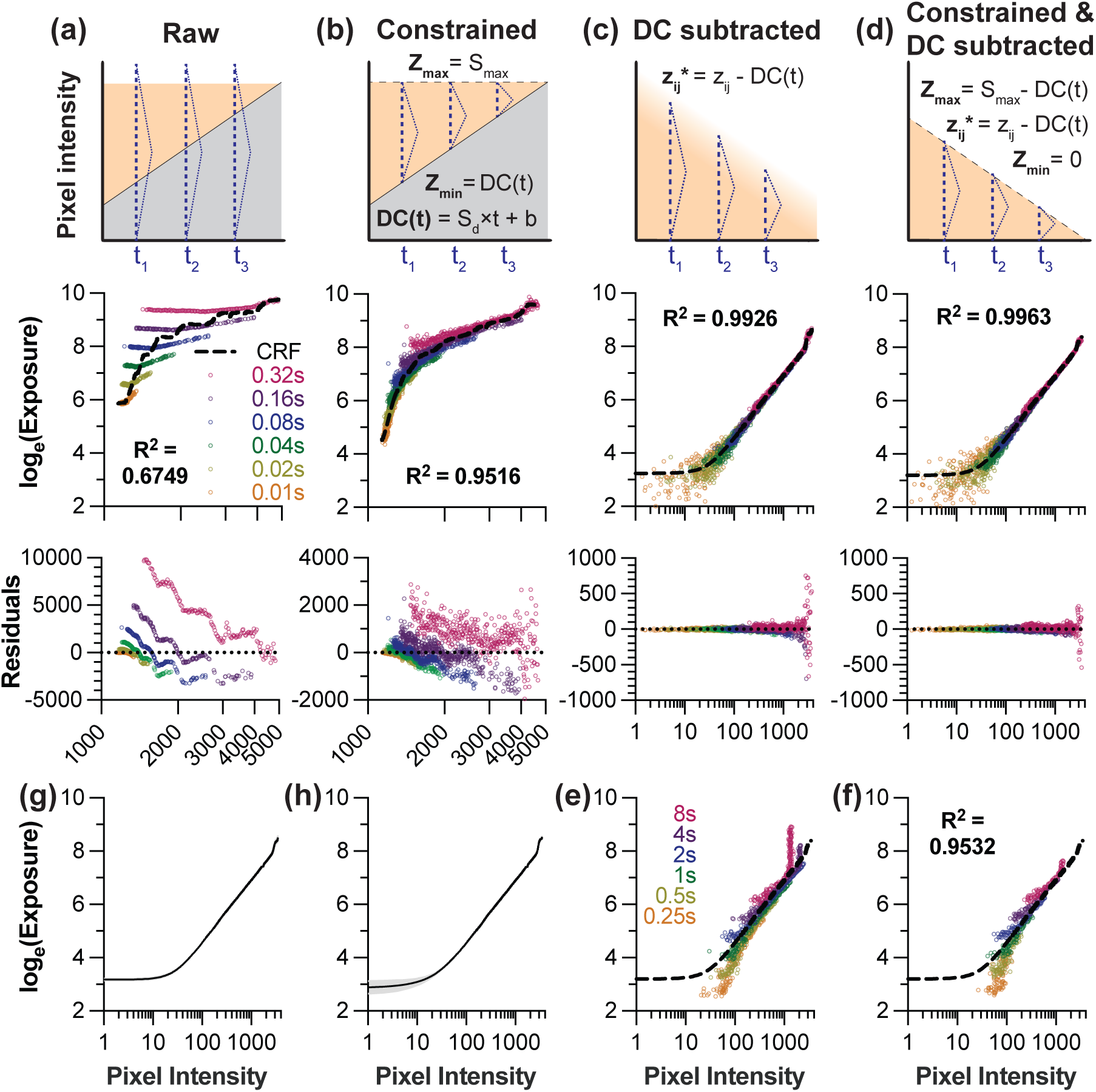
Signal preprocessing critically improves camera response function recovery in SWIR imaging. (a-d) Systematic evaluation of preprocessing strategies using exposures from 0.01-0.32 s. Each column shows the preprocessing schematic (top), resulting CRF fit with R² value (middle), and residual plot (bottom). In the schematics, the grey shading indicates the dark current noise (DC), orange shading represents the signal intensity range (*z_ij_*), and the purple triangle represents the Debevec weighting function *w*(*z_ij_*). (a) Raw intensity samples yield poor fits (R² = 0.675) with large and heteroscedastic residuals. (b) Constraining signals to characterized bounds (*Z_min_* to *Z_max_*) provides notable improvement (R² = 0.952). (c) DC subtraction alone substantially improves linearity (R² = 0.993) but retains outliers from unpredictable high-intensity signals. (d) Combining DC subtraction with capping the maximum pixel intensity yields optimal fits (R² = 0.996) with normally distributed residuals. (e-f) Impact of using long exposures (0.25-8 s) in CRF recovery. (e) In the absence of capping, saturated pixels from long exposure times contaminate the CRF across a broad pixel intensity range following denoising. (f) Combining noise subtraction with signal capping maintains reasonable congruence with the CRF from (d) (dashed line) with exposure times from 0.25 -8 s. (g) CRF consistency across different HDR weighting functions (Debevec’s triangle function, Robertson’s Gaussian function, Reinhard’s broadhat function, and Vinegoni’s window function).^18,20–22^ (h) CRF invariance across emission filters (2 LP and 6 BP filters). Black lines in (g-h) represent averaged CRFs, and gray shading represents the nominal standard deviation.

Without preprocessing, raw intensity samples produced poor CRF fits (R² = 0.68) with non-normally distributed residuals (Fig. 4a). The high DC and unpredictable behavior near saturation created systematic errors that violated the assumptions of classic HDR algorithms. We evaluated preprocessing strategies to address these issues. Capping pixel values at the characterized saturation point pixels (*z* > *S_max_* → *z*\* = *S_max_*, where *z\** represents the adjusted pixel intensity) provided modest improvement (data not shown, R^2^ = 0.73), preventing unpredictable high-intensity signals from corrupting the fit as our 16-bit InGaAs sensor exhibits unpredictable behavior well before the theoretical bit-depth limit. Further gains were achieved by additionally constraining sub-noise-floor pixels to *DC*(*t*) (*z* < *DC*(*t*) → *z\** = *DC*(*t*); R^2^ = 0.95, Fig. 4b), creating bounded signals within the characterized reliable range. However, DC subtraction was the most impactful preprocessing step (*z* < *DC*(*t*) → *z\** = *0*; *Z_min_* = 0*; z\** = *z* – *DC*(*t*)), as it linearizes the relationship between exposure time and measured intensity to yield excellent CRF fits (R^2^ = 0.993) with normally distributed residuals (Fig. 4c). Combining both preprocessing steps—subtracting DC and constraining the saturation limit—yielded further improved CRF fits (R² > 0.996) suitable for quantitative imaging (Fig. 4d).

The combination of DC subtraction and signal capping proved particularly critical when including longer exposure times in the CRF recovery. As exposure time increases, the growing DC floor progressively narrows the usable dynamic range, causing unstable near-saturation signals to appear at lower absolute pixel values. Without capping, these erratic signals introduce systematic errors across a broader range of the CRF (Fig. 4e), meaning that the denoised data cannot be effectively used for longer exposure times without capping the pixel intensities at *S_max_*. In contrast, proper signal capping enables fit quality to be maintained even with exposures up to 8 s (Fig. 4f). The consistency in the CRF between Fig 4d and 4f confirmed that CRFs recovered using shorter exposures (0.01-0.32s) can be reliably applied to all exposure times, eliminating the need to include noise-dominated long exposures in the calibration process. More specifically, the CRF recovered with short exposures fit the samples acquired at long exposures (R^2^ = 0.9532) with similarly distributed residuals, indicating that the short-exposure CRF be used to recover intensities obtained at longer exposures.

Selection of the pixel weighting approach from the list of available options could affect CRF recovery and image fusion. Vinegoni used the simplest weighting to improve algorithm speed for real-time, confocal or two-photon imaging settings: a window function ranging between the detector noise floor and saturation.^20^ Debevec’s triangular weighting provides smoother transitions at the bounds, reducing edge artifacts in the CRF recovery.^18^ Alternatively, Mitsunaga and Nayar use a weighting function that is a derivative of the response function itself. Such response function was recovered using an alternate method involving estimating irradiance.^23^ This function emphasizes higher pixel values because they tend to have higher SNR. Reinhard modified this method by multiplying the weighting function by a broadhat function that forces weights near the minimum and maximum to approach 0.^21,24^ Robertson also proposed weighting pixels at longer exposures and fitting the final camera response to a cubic spline curve while using a gaussian weighting funciton.^22^ We performed CRF recovery with each of the weighting functions discussed (data not shown). As long as the function weights intensities near the noise floor and saturation point close to 0, the correlation coefficient for the CRF fit on the intensity samples was above 0.99. This applied for the weighting functions from Vinegoni, Reinhard (broadhat), Robertson, and Debevec.^18,20,22,24^ The CRFs recovered with these four weighting functions were congruent (Fig. 3g). We selected Debevec’s method for subsequent calculations as it provides the best balance of computational efficiency and accuracy for our offline processing workflow.

In classical HDR image processing with RGB cameras, CRF recovery is repeated for each color channel, as each channel has a distinct, frequency-dependent light response.^18,25^ We did not observe such differences when recovering the CRF using intensities sampled from different combinations of excitation lasers and filter sets (Fig. 4h) and thus proceeded using the same CRF for each of the bandpass filters and two longpass filters.

The invariance of the recovered CRF across different weighting functions and emission filters significantly streamlines the HDR workflow, as the robustness of the CRF implies that users need only perform CRF recovery once for the detector/imaging system rather than for each experimental configuration.

### 3.3 HDR Imaging Extends Dynamic Range for Quantitative Fluorophore Detection

To validate the HDR method for quantitative fluorescence imaging across wide contrast agent concentration ranges, we imaged serial dilutions of ICG *in vitro*. This experiment directly addresses a practical challenge in preclinical imaging studies: simultaneously quantifying contrast agents that accumulate at vastly different concentrations across tissues.

We prepared a 1:2 dilution series of ICG (6 μM to 0.003 μM) and imaged with exposures ranging from 0.01 to 10.24 s. The 0.01 s exposure was considered underexposed because none of the features reached >20% of *Z_max_*, and the 10.24 s exposure was considered overexposed because most of the average well intensities were at or above *Z_max._* Therefore, only exposures between 0.02 s and 5.12 s were used in image fusion. Single-exposure imaging revealed a tradeoff (Fig. 5a-b): a short exposure (0.02 s) captured the highest concentration (6 µM) without saturation but failed to detect ICG below 0.012 μM, while longer exposures (0.32 s) revealed the dimmest wells (0.003 µM) but saturated at concentrations above 0.375 μM. The single exposure time images could resolve approximately 2-2.5 orders of magnitude of concentrations (Fig. 5c), but with increasing uncertainty at the lower pixel intensity values (Fig. 5b).

**Figure 5.**
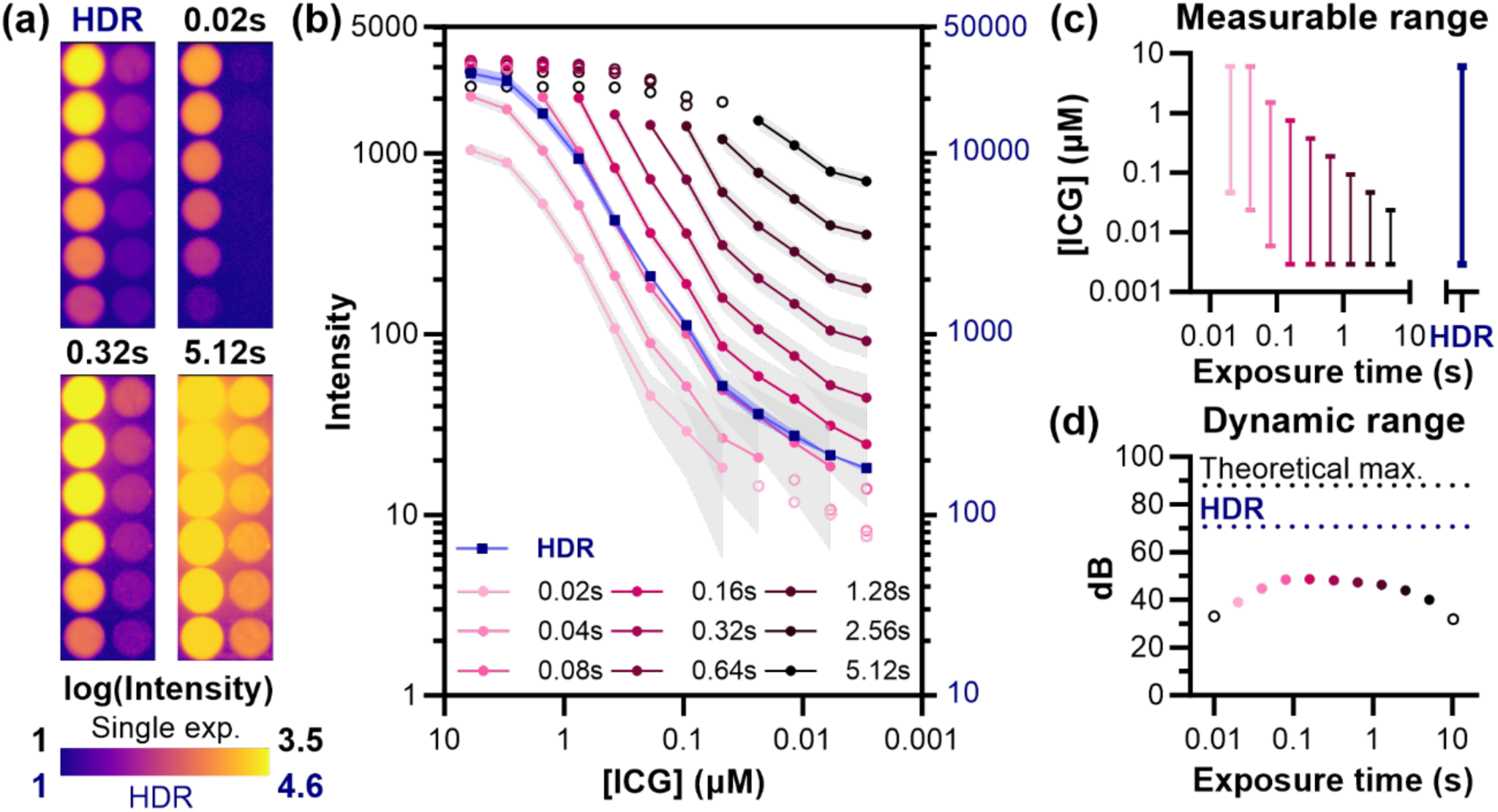
HDR imaging enables quantification across three orders of magnitude of fluorophore concentration. (a) ICG dilution series (1:2, starting at 6 μM) imaged with single exposures using a 1000 nm NIR-II LP filter and HDR fusion, displayed on log₁₀ scale. (b) Quantitative analysis of mean ICG intensity (±SD) per well across exposure times. Unfilled markers indicate measurements outside the reliable range (below noise floor or above saturation). (c) Measurable concentration range for each exposure time, demonstrating that no single exposure captures the full dilution series. (d) Dynamic range comparison showing a 22 dB improvement with HDR fusion (70.2 dB) over the best single exposure (48.2 dB). The DR of the HDR image is limited by contrast agent dilution rather than the instrumentation; the theoretical maximum HDR DR for this exposure time range (88 dB) is shown as well. Unfilled, black markers indicate exposure times (0.01, 10.24 s) not shown in panel (b).

HDR fusion image, on the other hand, simultaneously captured all ICG concentrations across the three-order-of-magnitude range, achieving a dynamic range of 70.2 dB—a 22 dB improvement over the best single exposure (Fig. 4d). Higher exposures for single images have limited dynamic range because of the increasing DC floor and variance, which places a limit on viable exposure times equivalent to where DC saturates pixels fully. With HDR imaging, the highest measurable value is dependent on the highest measured signal at the lowest exposure possible. Using Equation 9, the theoretical maximum dynamic range in this imaging system using HDR image fusion for the exposure time range shown to be useful for this ICG dilution (0.02 – 5.12 s) would be ∼88 dB, assuming pixels reach saturation at the lowest viable exposure time of 0.02s. The self-quenching of ICG at the highest concentration (6 µM) and shortest exposure time (0.02s) evident in the signal intensity plateau seen for this exposure time prevents the capture of this maximal HDR range in this particular data set. Overall, the dynamic range improvement allows us to be limited by contrast agent brightness and administration method and not by our imaging system.^26^

### 3.4 HDR Imaging Improves Contrast and Quantitative Output in Preclinical Imaging

Seeing the improvement of dynamic range in SWIR fluorophore imaging, we investigated how this improvement translates to *in vivo* imaging. Imaging of nanoparticles after administration to small animal models is of particular interest because particles ≥6 nm in diameter tend to accumulate in the liver,^27,28^ resulting in extremely bright abdominal fluorescence. To enable visualization of features other than the liver, one may need to cover the abdominal region to avoid image saturation. Because signals from these mouse models can have a wide intensity range, HDR image processing is advantageous for ensuring the signal is neither noisy nor saturated.^26^

Figure 6 demonstrates this challenge with images of SWIR-emitting quantum dots 1 hr post-injection. Single exposures force a compromise: short exposures (0.04 s) maintain liver signal within the quantifiable range but compress vascular signals near the noise floor, reducing measurement precision. Longer exposures (0.32 s) spread vascular intensities across a wider range for better quantification but have saturated pixels in the liver. Critically, selecting the “optimal” exposure requires prior knowledge of fluorophore biodistribution and brightness, which is challenging in an experimental setting.

**Figure 6.**
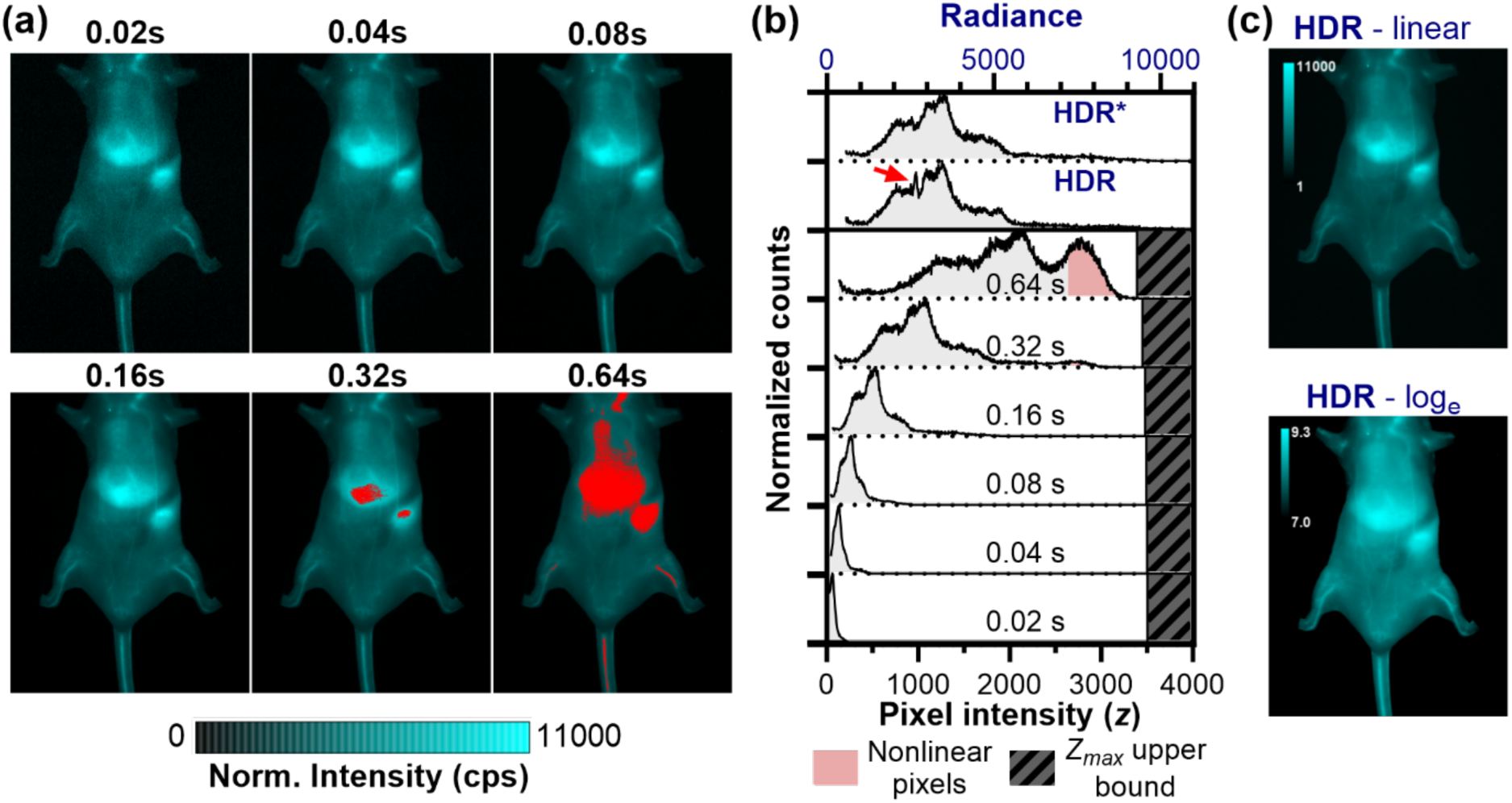
HDR imaging overcomes the dynamic range limitations of single-exposure SWIR imaging. (a) Exposure time normalized images of a mouse injected with PbS/CdS quantum dots (5 mg/kg i.v.) at 1-hr post-injection, imaged with 808 nm excitation and 1300 nm bandpass filter at multiple exposures between 0.02 and 0.64 s. Red regions indicate saturated pixels. (b) HDR radiance histograms and pixel intensity histograms from single-exposure time images with near-saturation pixels exhibiting non-linear behavior shaded red. Artifact in HDR radiance histogram (red arrow) is eliminated by excluding repeated near-noise (<5% of *Z_max_*) and near-saturation exposures (>95% of *Z_max_*) on a per-pixel basis (HDR*). (c) Linear-scale and log-scale visualization of the HDR image.

HDR imaging eliminates this guesswork. By acquiring multiple exposures systematically, the method captures all features across their optimal intensity ranges, then fuses them into a single quantitatively accurate image. The HDR radiance histogram stretches the radiance values over a wide range, enabling more nuanced quantitative analysis compared to single exposure time measurements. The linear-scale HDR image preserves the quantitative relationships needed for biodistribution analysis, while log-scale visualization reveals dimmer anatomical details typically masked by the comparatively bright signal in the liver. Though showing images on a linear scale is typical for presentation of scientific images, displaying the HDR image on a log-scale is an alternative that is much closer to how humans perceive light intensity. Practically, the log-scale image emphasizes low-intensity features such as small vessels and can be advantageous for qualitative viewing.

We occasionally observe artifacts in the HDR radiance histogram, which we attribute to over-usage of high-variance, non-linear intensities near *Z_max_* and *Z_min_*. One such artifact is displayed in the HDR histogram in Fig. 6b (red arrow). Approaches to limit variance increases with image fusion have been described, such as though eliminating weak, noisy exposures from image fusion and removing overexposed parts of images entirely,^16^ and prioritizing high SNR pixels.^20,29^ We similarly adapt the HDR fusion algorithm to avoid the inclusion of multiple near-noise or near-saturation exposures. Specifically, if an exposure yielded a pixel intensity at the upper bounds near *Z_max_* (i.e., *z*\* > 0.95*Z_max_*), then longer exposure times were excluded from the HDR fusion for that pixel. Likewise, if an exposure yielded a very low pixel intensity close to *Z_min_* (defined as *z*\* < 0.05*Z_max_* as *Z_min_* is 0 following DC subtraction), shorter exposure times were excluded from the HDR fusion for that pixel to limit the unnecessary inclusion of noise in the image. This approach ensured that at least one exposure time is available for all pixels, even when the brightest pixel intensity is very close to the noise floor or the dimmest pixel intensity is very close to the saturation limit, while limiting the noise or artifacts introduced by including additional pixels that do not contain useful information. This modification successfully eliminated the artifact; the resulting HDR image histogram profile closely resembled that of the longer, non-saturated single exposures. To quantify the practical benefits of HDR imaging for small animal imaging, we analyzed contrast-to-noise ratios (CNR) across key anatomical features. Figure 7 shows line scans drawn over organs and vessels for representative mice administered either ICG (Fig. 7a) or PbS/CdS QDs (Fig. 7b). At short exposure times, vessels are poorly exposed, but the liver exhibits optimal single-exposure CNR. In contrast, the longest exposure time best highlights signal from low-intensity features (typically vasculature). Overall, the exposure time that maximizes CNR for each pixel in the line profile varies. After HDR image fusion, all features can be visualized simultaneously in a single image without any saturated pixels. Moreover, this exposure-adjusted HDR image can be used in downstream fluorophore quantitation without needing to consider exposure time, and radiance maps acquired using the same camera and CRF can be compared between each other without normalization for imaging conditions except illumination intensity.^18^ We also observed a consistent improvement in dynamic range, typically ∼10 dB in preclinical imaging. HDR fusion eliminates feature-dependent optimization, presenting each anatomical structure at its ideal contrast. These improvements translate directly to practical benefits: researchers can quantify biodistribution across all organs from a single dataset without prior knowledge of optimal imaging parameters, streamlining experimental workflows while ensuring no anatomical features are sacrificed to technical limitations.

**Fig. 7.**
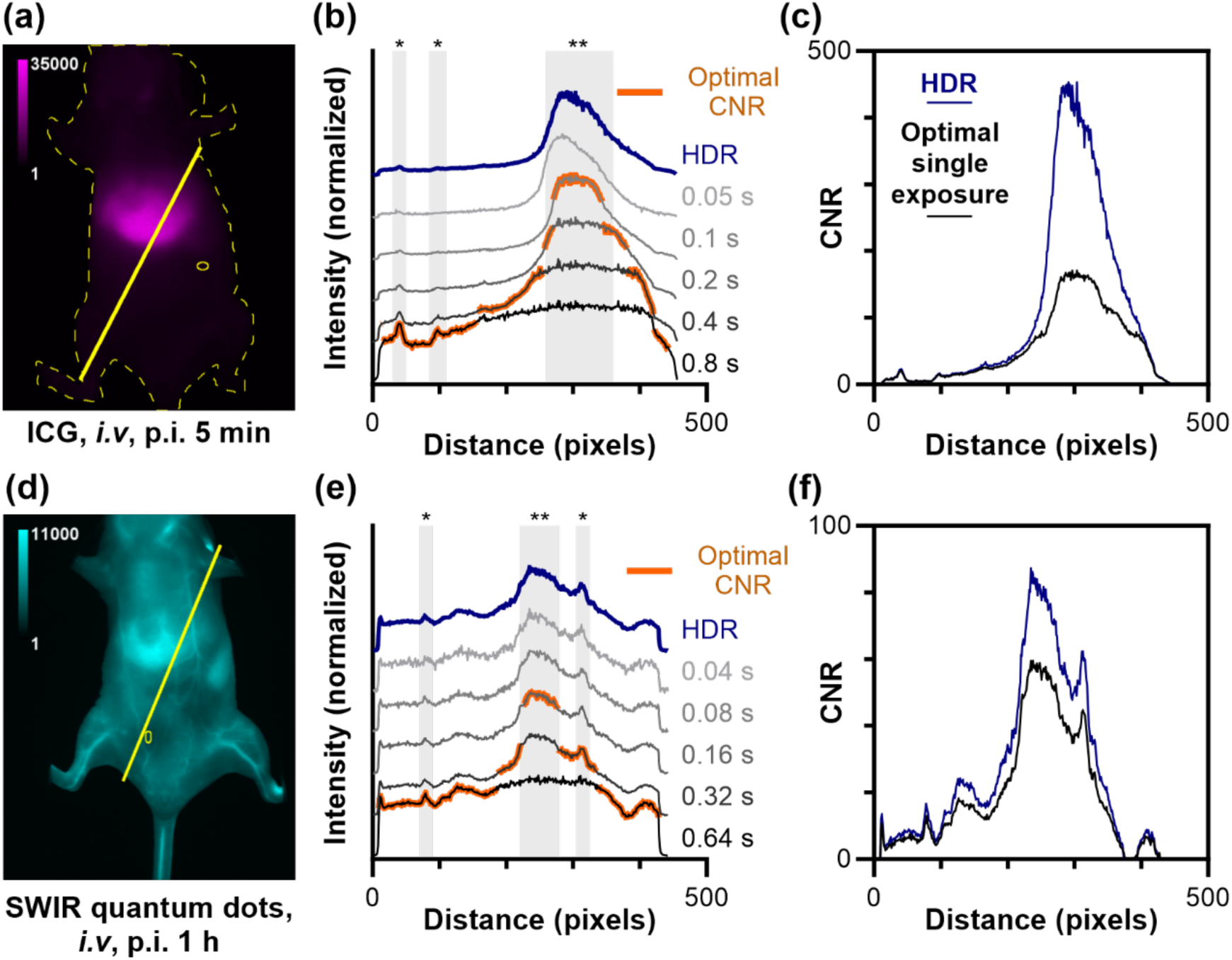
Representative HDR images of mouse. administered (a) ICG (imaged with 1150 BP filter) and (d) SWIR-emitting PbS/CdS QDs (imaged with 1300 BP filter), 1-hour post-injection. Line profiles (solid yellow line) are drawn through the liver and across the surrounding tissue; yellow oval indicates region used for on-mouse background signal. (b, e) Line profile regions highlighted in orange have the highest CNR of the single exposure images, which varies with region. Liver (**) and blood vessel (*) features are indicated by gray-shaded regions.(c, f) Comparison of the CNR for a single exposure and HDR images.

## 4 Conclusions

We successfully developed an HDR method for SWIR imaging that addresses the fundamental dynamic range limitations of InGaAs cameras in preclinical fluorescence imaging applications. Through careful adaptation of classical HDR algorithms to incorporate exposure-dependent dark current correction, signal preprocessing and dynamic weighting, we achieve a >22 dB improvement in dynamic range over single-exposure imaging. This advancement enables quantitative fluorescence analysis over more than three orders of magnitude of contrast agent concentration from a single acquisition sequence. For preclinical imaging applications, this eliminates the trade-off between capturing bright organs (liver, spleen) and dim vascular features, streamlining experimental workflows while preserving quantitative accuracy across anatomical structures. The practical application of this method requires only software processing and standard imaging protocols, making it accessible to existing SWIR imaging systems without hardware modifications. One-time camera characterization generates calibration models that enable rapid processing of subsequent datasets. By extending quantitative imaging capabilities across the full range of anatomical brightness levels, this HDR approach unlocks the full potential of SWIR fluorescence imaging for biodistribution studies and preclinical research.

## Code and Data Availability

All data presented in this article are available upon reasonable request. The code utilized for image processing available on GitHub at [link will be inserted before publication]. Plotting of data was performed using GraphPad Prism 10 and figure layouts were prepared in GraphPad Prism 10 and Adobe Illustrator CC 2024. Claude by Anthropic was used for modification of HDR algorithm script to enable integration into image processing workflow.

## Acknowledgments

The research reported in this publication was supported by the National Institute of General Medical Sciences (NIGMS, Award No. R01GM129437) and the National Institute of Biomedical Imaging and Bioengineering (NIBIB, Award No. R21EB032647) of the National Institutes of Health (NIH). Claude Sonnet 4 was used during manuscript editing for summarization and to improve language clarity.

